# Genetic Developmental Timing Revealed by Inter-Species Transplantations in Fish

**DOI:** 10.1101/2020.04.02.019471

**Authors:** Jana Franziska Fuhrmann, Lorena Buono, Juan Ramón Martinez Morales, Lázaro Centanin

## Abstract

The path from a fertilised egg to an embryo involves the coordinated formation of cell types, tissues and organs. Developmental modules (Raff, 1996) comprise discrete units specified by self-sufficient genetic programs that can interact among each other during embryogenesis. Here we took advantage of the different span of embryonic development between two far related teleosts, zebrafish (*Danio rerio*) and medaka (*Oryzias latipes*), of 3 and 9 days respectively, to explore modularity principles. We report that inter-species blastula transplantations result in the ectopic formation of a retina formed by donor cells — a module. We show that the developmental time of the retina follows a genetic program: an ectopic zebrafish retina in medaka develops with zebrafish dynamics. Heterologous transplantation results in a temporal decoupling between the donor retina and host organism, illustrated by two paradigms that require retina-host interactions: lens recruitment and retino-tectal projections. Our results uncover a new experimental system to address temporal decoupling along embryonic development, and highlight the presence of largely autonomous but yet interconnected developmental modules orchestrating organogenesis.

## Introduction

In vertebrates, organogenesis takes place during embryonic development and follows a stereotypic, species-specific timing. Cases in which two organs of the same type are generated within an organism — eyes, ears, lungs, kidneys, gonads — indicate that these develop in a synchronized manner despite constituting independent units. The temporal control of organogenesis is of paramount importance to secure a functional coordination of organs within systems, i.e. neurons in a sensory organ should mature and become functional together with their target regions. A long-standing question in the field is whether neural organs follow an endogenous timing that defines the onset of neurogenesis — the autonomous timing —, or if alternatively, there are global signals that guarantee coordination among cell types, tissues, and organs — the ontogenic timing.

The vertebrate neural retina constitutes a major model for neurogenesis in the central nervous system (CNC), and it has been long ago demonstrated that the different types of retinal neurons are formed in a stereotypic temporal order, and arranged in dedicated layers [1]. Retinal organoids and aggregates demonstrated that the vertebrate retina is capable of autonomously patterning [2], earlier suggested by the transplantation of optic vesicles into ectopic regions of the chick and fish embryo [3]. Although the self-organizing properties of the neural retina were demonstrated for other organoids, the inherently artificial conditions to develop 3D cultures, or the technical artefacts that accompany the transplantation of an optic cup, might affect the temporal sequence of biological processes on organoids or aggregates [4]. The ideal set up to explore developmental timing, therefore, should exploit the self-organizing properties of the retina while developing in a homo – or heterochronic physiological environment.

Teleost fish represent the vertebrate clade with most species, which span across a huge range of sizes, body shapes and times of embryonic development [5]. Among teleost fish, *Danio rerio* and *Oryzias latipes* (zebrafish and medaka, respectively) have diverged some 250 million years ago, and belong to two of the most distant subgroups [6]. One of the most obvious differences between them is the time that they take to complete embryonic development, 3 days for zebrafish and 9 days for medaka (Figure 1A). Here, we use zebrafish - medaka inter-species transplantation to report that isochronic transplantation of blastomeres from one species into the other results in the formation of an ectopic retina formed by donor cells. We exploited this unique set-up to study the development of an entire organ on an extrinsic, yet physiological, environment. Using fluorescent transgenic report lines and RNA-seq analysis, we show that the ectopic retina develops with a genetic timing, irrespective of the host.

**Figure 1.**
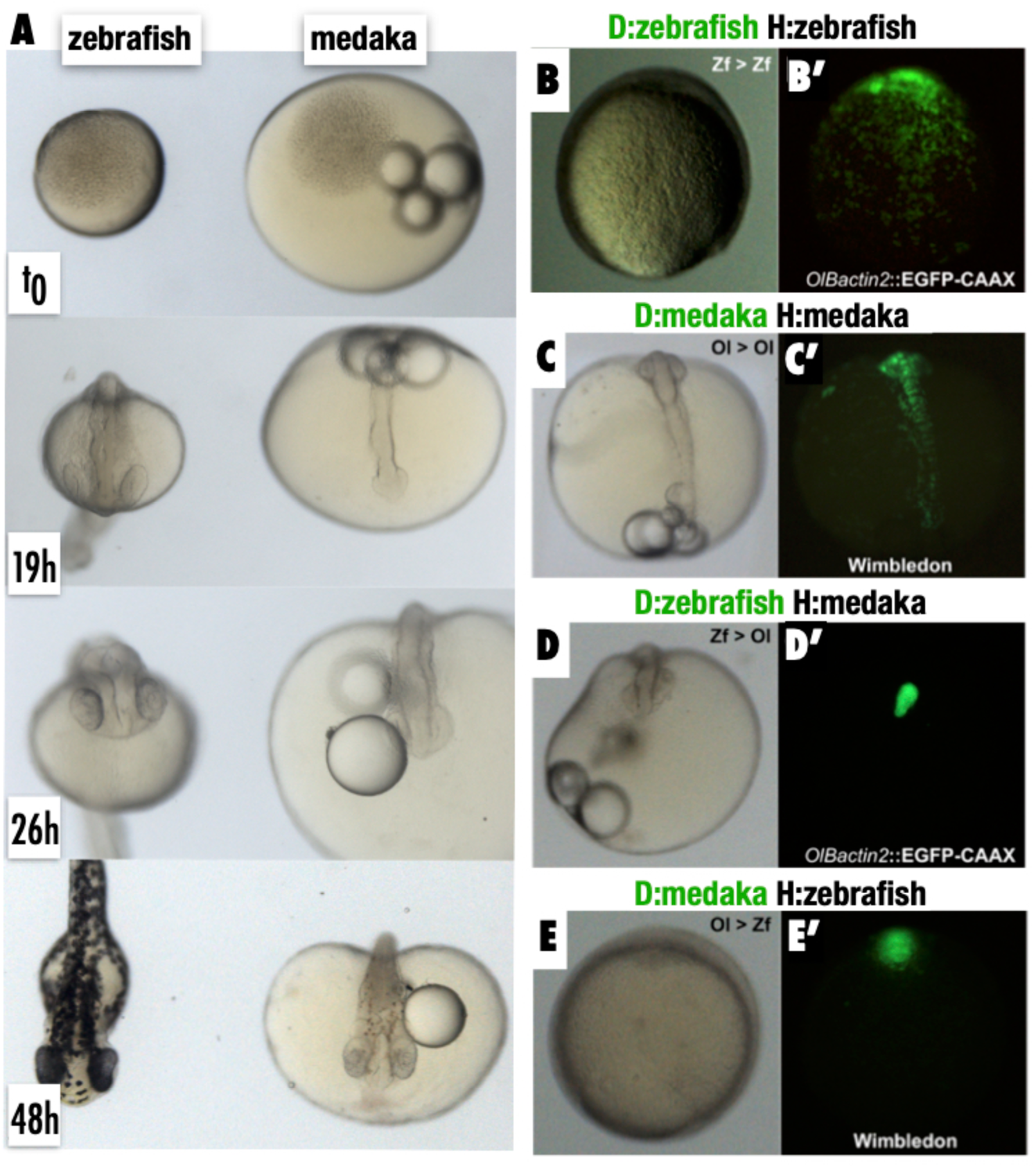
Intra and Inter-species transplantation of blastocysts in zebrafish and medaka. **(A)** Binocular images of zebrafish (left) and medaka (right) embryos at different stages of early embryonic development, synchronised at the blastula stage (512 cells). Notice that zebrafish develops faster that medaka. **(B - E)** Transmitted and fluorescent images of non-labeled embryos transplanted with isochronic EGFP+ blastocysts. Intraspecies transplantations results in EGFP+ cells intermingling with the host cells both in zebrafish (B-B’) and medaka (C, C’). Inter-species transplantations result in a different scenario, where donor cells stay cluster together both in zebraka (D, D’) and in medrafish (E, E’).

## Results & Discussion

### Medaka / zebrafish inter-species transplantation results in the formation of ectopic retinae

Genetic chimeras formed by transplanting blastomeres from one fish embryo to another have been extensively used in zebrafish (Figure 1A, left) and medaka (Figure 1A, right), i.e to define the cell-autonomous vs cell-non-autonomous roles of novel mutant lines, to study lineages during embryonic and post embryonic development, *etc*. [7-11]. Intra-species transplantations at blastula stage results in donor and host cells mixed along the developing embryo. This was indeed the case when we transplanted blastomeres from ubiquitously labelled transgenic lines (*ßactin*::CAAX-EYFP in zebrafish, *Wimbledon* or *Gaudi*^*LoxP*.*OUT*^ in medaka)[7,12] into non-labelled controls from the same species (Figure 1B,C’). We have observed that in trans-species transplantations, however, donor cells stay clustered together and do not mix with host cells during gastrulation, both for <u>zebra</u>fish-to-med<u>aka</u> (zebraka, Figure 1D,D’) and for <u>med</u>aka-to-zeb<u>rafish</u> (medrafish) (Figure1E, E’) (N>100 chimeras for each case). In both species, host blastomeres can proceed through gastrulation and a body axis is evident at 19 hours post-transplantation (hpt) (Figure 1D). Stunningly, the cluster of transplanted cells often develops into an ectopic organ that resembled a retina, which is formed exclusively by EGFP-positive cells (23 zebraka embryos out of 72 transplanted, *Experiment 106*) (Figure 2A). We have observed ectopic retinas for both trans-species transplantation setups, often containing retinal pigmented epithelium (Figure 2B), which indicates that the initial alien cluster followed a developmental program despite of not intermingling with host cells during epiboly. The expression of retinal transcripts in the cluster was confirmed by using transcriptional reporters for retinal progenitors and retinal neurons in medaka, namely *Rx2* - *retinal homeobox factor 2* [11,13,14] and a*toh7* [9,15-17]. When blastomeres from medaka Tg(*Rx2*:H2B-RFP) were transplanted into non-labelled zebrafish blastulae, the retinal-like cluster expressed the medaka retinal reporter *Rx2* (Figure 2C). The same result has been obtained when we used Tg(*atoh7*:EGFP) as donors, suggesting that the ectopic cluster has both retinal identity and the potential to trigger neurogenesis (Figure 2D).

**Figure 2.**
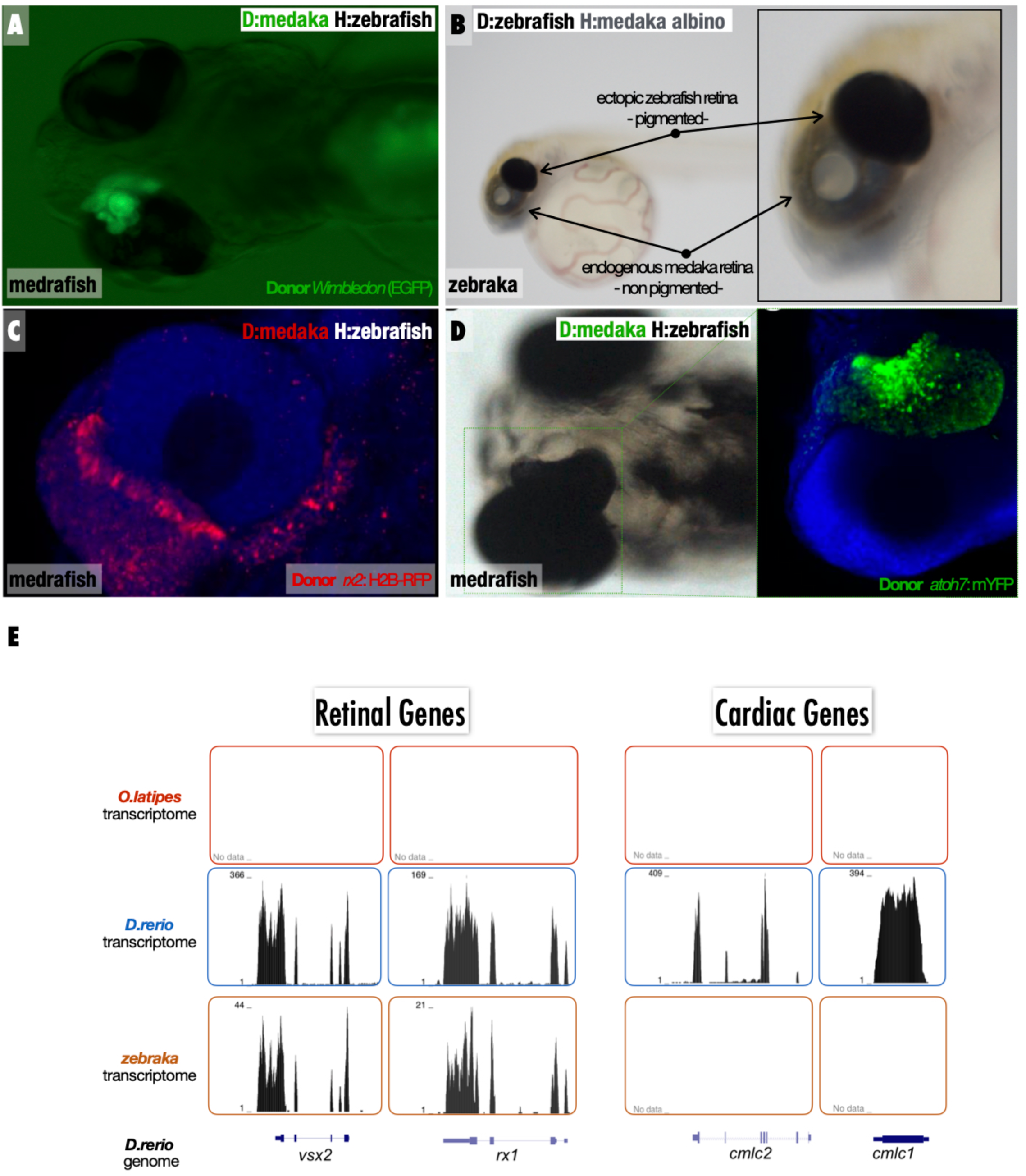
EGFP+ cluster develops into an ectopic retina both in zebraka and medrafish. **(A-B)** Binocular and **(C-D)** confocal images of medrafish (A, C, D) and zebraka (B). The EGFP+ (A) or pigmented (B) cluster develops into an ectopic retina, which expresses retinal progenitor (Rx2:H2B-RFP donors, C) and neurogenic (*Atoh7*:mYFP donors, D) markers. **(E)** Transcriptomes of medaka (upper), zebrafish (middle) and zebraka (bottom) plotted along the zebrafish genome. The zebrafish cells in medrafish display retinal identity (*vsx2, rx1*, left two panels) and no cardiac genes markers (*cmlc1&2*, right panels).

There are a number of methods available to address species-specific contribution of transcripts to chimera [18]. The low sequence identity of homologous genes between zebrafish and medaka allows segregating the chimeric transcriptome *in silico*, therefore expanding the analysis of the transcriptional profile in ectopic retinae of zebrakas. We compare RNA-seq from zebrafish, medaka, and zebraka embryos at 48 hpf, a stage in which neurogenesis has started in zebrafish but it is only about to start in medaka. We selected chimeras with a clear EGFP+ cluster close to one of the endogenous retinae, as these are the most likely to become a retina. As expected, the transcriptome from zebrafish, but not the one from medaka, aligned to the zebrafish genome (Figure 2E, Supplementary Figure 1). Due to the large evolutionary distance, only a few RNA-seq reads form the medaka transcriptome mapped on the zebrafish genome (Supplementary figure 1), and these displayed a distinguishable morphology as well as a minimal number of reads (Supplementary Figure 1). In contrast, numerous reads from the transcriptome of zebrakas, consisting in a full medaka transcriptome plus a partial transcriptome from the few zebrafish cells, could be aligned to the zebrafish genome (Figure 2E, Supplementary Figure 1). These peaks corresponded to genes that are expressed by the zebrafish retina at the same developmental time (*vsx2, rx1, rx2* among others, Figure 2E and Table 1). Genes exclusively expressed in other organs at the same stage, and therefore present in the zebrafish transcriptome (i.e., cardiac muscle) were absent in the zebraka transcriptome (Figure 2E). This analysis confirms the retinal identity of the EGFP+ cluster in zebrakas chimeras.

**Table 1.**
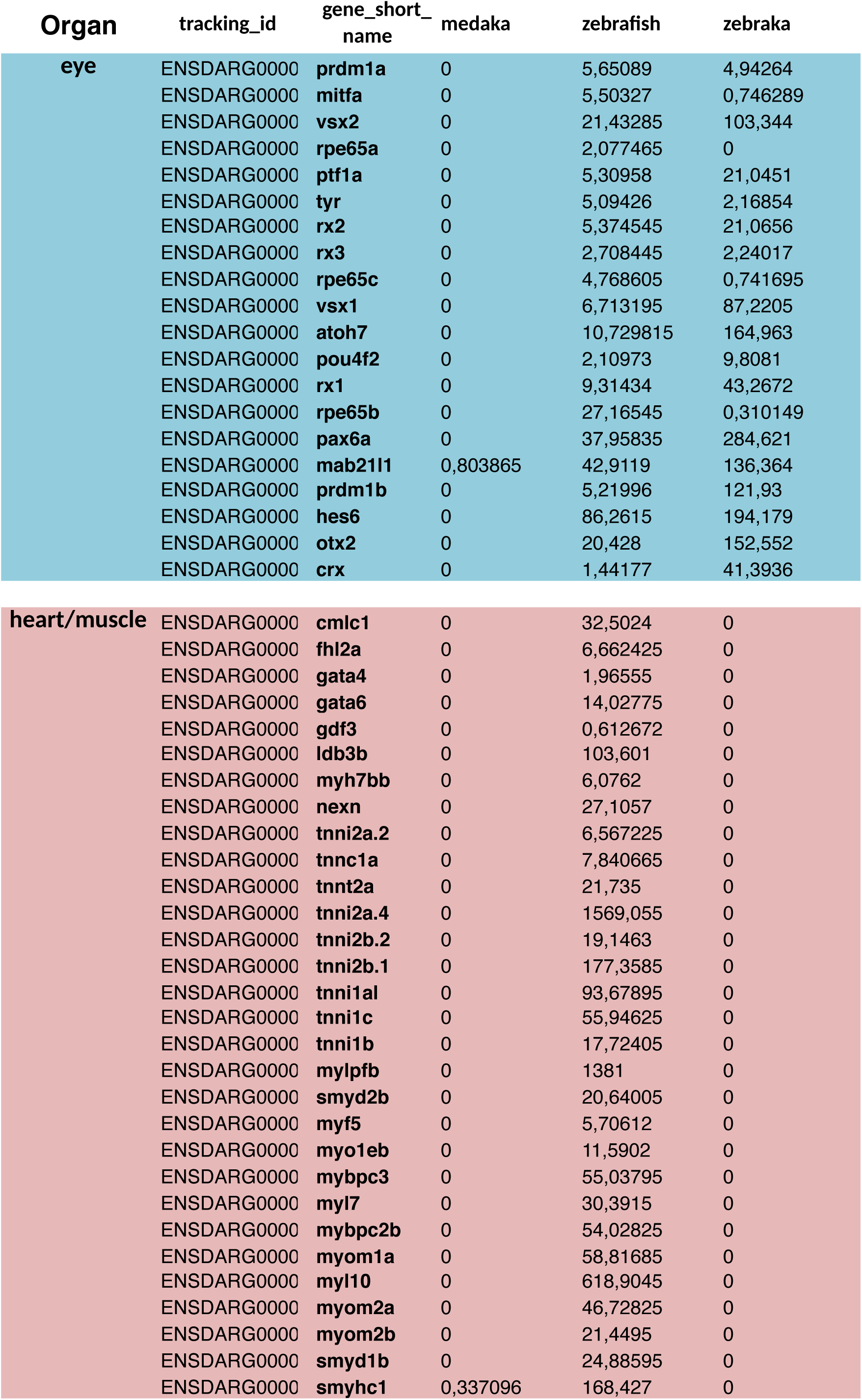
List of zebrafish transcripts (and their ID) corresponding to retina genes (upper section) and heart genes (bottom section). The different columns show the count number of their respective sequences in transcriptomes from medaka, zebrafish and zebraka.

It has long been proposed that neural ectoderm constitutes the default differentiation program for early embryonic cells. Indeed, neural retina was the first *organoid* produced in 3D cultures from an aggregate of embryonic stem cells (ESCs)[2], followed by other neural organs [19,20]. In zebraka and medrafish chimeras, we have noticed that ectopic retinae inevitably form adjacent to an endogenous retina (Figure 2A, B, N>60 chimerae). We have never observed an ectopic retina in remote locations, which strongly suggests that positional information from the host can be decoded by the ectopic cells - although does not elucidate whether the host anlage has a permissive or an inductive role. Retinal identity is not the only fate that ectopic blastocysts can adopt when transplanted into the foreign species, since we have observed clusters differentiating into vasculature or pigmented cells (Supplementary Figure 2) [21]. The ectopic retina obtained in the present inter-species transplantation protocol, however, is the only organ entirely composed of foreign cells (Figure 2A) and as such, permits a compartmentalised analysis of host and donor organs going through embryonic development in parallel and in the same embryonic environment.

### Ectopic retinae differentiation follows a genetic timing

The onset of retinal neurogenesis in medaka and zebrafish occurs at different hours post-fertilization (hpf). We decided to use the inter-species chimeras as a paradigm to address developmental timing, i.e., to explore whether retinal neurogenesis follows an intrinsic temporal program (*genetic* timing) or if alternatively, it responds to signals from neighbour tissues operating as temporal coordinators (*ontogenic* timing). Using the transcriptome data that we obtained from zebrakas at 48 hpf, we aimed at analysing the relative expression of progenitor and neurogenic genes in the ectopic retinae. We used *rx3* and *vsx2* as retinal progenitor markers and *atoh7, ptf1a* and *pou42* as neurogenic/differentiation markers [22-25], and compared their expression ratios in zebrafish, medaka and zebrakas, using as a scaffold published transcriptomes from zebrafish and medaka at different developmental stages [26]. We noticed that ratios in the zebrafish transcriptome of zebrakas match those of wild type zebrafish, while ratios in the medaka transcriptome of zebraka match those of wild type medakas (Supplementary Figure 3). These results indicate that despite developing in a foreign species, a zebrafish retina in zebraka keeps its genetic differentiation dynamics.

To confirm the genetic timing of retinogenesis in zebrakas *in vivo*, we performed transplantations using a Tg(*atoh7*:EGFP) zebrafish as a donor. Retinal ganglion cells (RGCs) represent the first cell type to differentiate in the vertebrate neural retina, and *atoh7* has extensively been used as a marker to follow the onset of RGCs in both zebrafish and medaka [9,15-17]. Atoh7 expression, and expression of fluorescent proteins under the control of an *atoh7* promoter, can be detected at 26 hpf in zebrafish and one day later (50hpf) in medaka. We transplanted transgenic Tg(*atoh7*:EGFP)(*ef1*:LynTomato) zebrafish blastocysts into unlabelled medaka blastula and observed the onset of EGFP expression by 26 hpf, well before the expression of the endogenous medaka *atoh7* (Figure 3A). This result complements our transcriptome analysis and indicate once again that RGC generation follows a genetic timing irrespective of the timing of their neighbouring cells.

**Figure 3.**
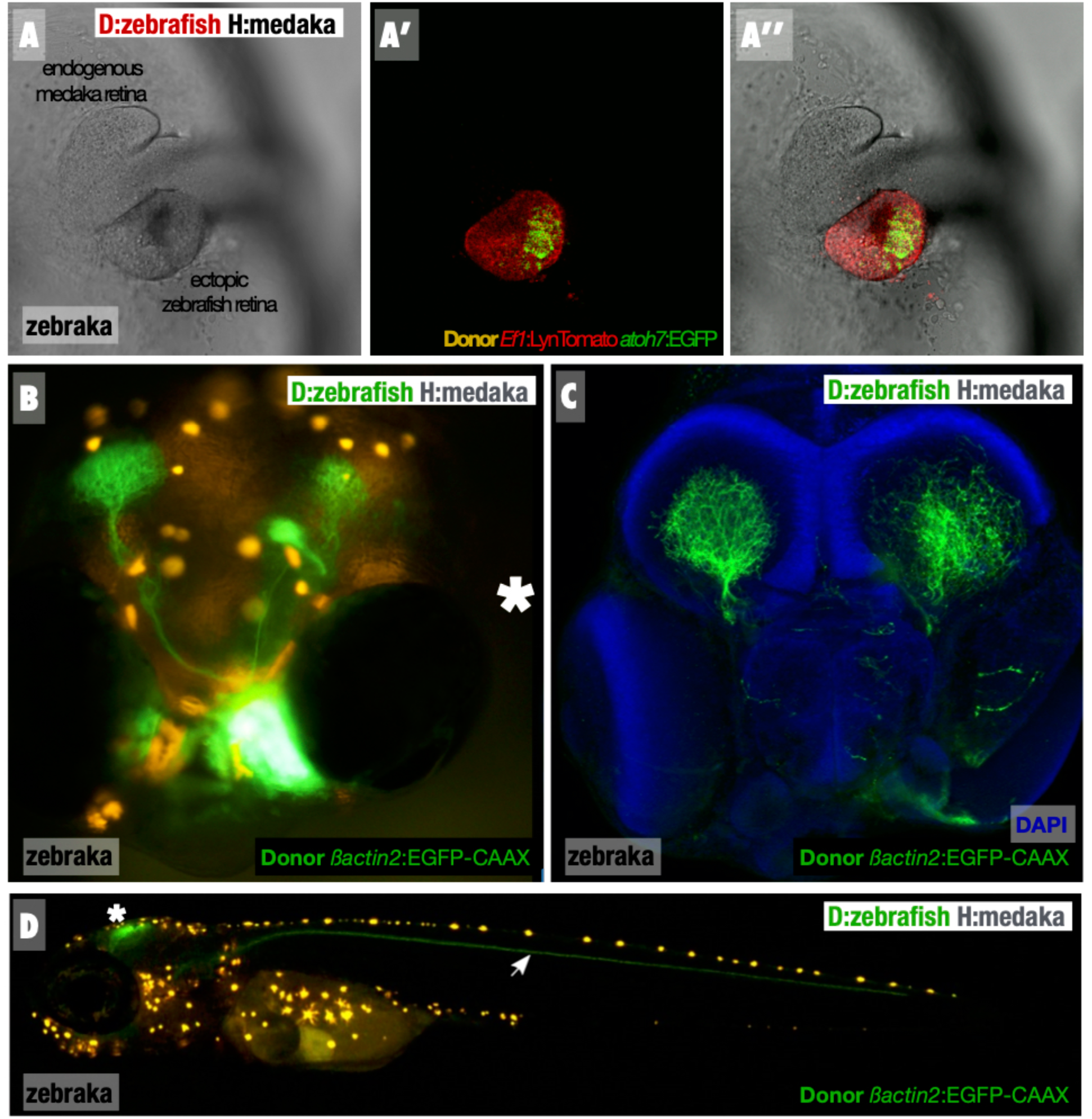
Retinogenesis follow a genetic developmental time. **(A-A’’)** Confocal images of a zebraka, where the donor is Tg(*Ef1*:LynTomato, *atoh7*:EGFP). EGFP expression in the zebrafish cluster is evident at the vesicle stage of the medaka host. **(B-C)** Binocular (B) and confocal (C) picture of a 6 dpf zebraka where the donor is Tg(*Olßactin2*:EGFP-CAAX). Note that the ectopic retina projects to the host tecti. Embryo is the same in both panels, imaged alive (B) or after fixation and staining (C). **(D)** Confocal image of a zebraka at 9dfp. A nerve coming from the ectopic zebrafish retina navigates along the posterior lateral line nerve. Other projections from the ectopic retina, presumably from later RGCs, project to the tectum.

### Excessive arborisation from zebrafish RGCs in the medaka optic tectum

The generation of premature RGCs by the ectopic EGFP+ cluster has consequences in the host that we revealed by analysing the optic nerve in zebrakas and medrafish. In vertebrates, RGCs projections group in an optic nerve that migrate from the retinae to their target region in the brain, the visual cortex in mammals and the optic tectum in fish. In zebrafish and medaka, each retina project an optic nerve to the contralateral optic tectum [27,28]. We noticed that in zebrakas where donor cells were Tg(*atoh7*:EGFP), zebrafish RGCs form an optic nerve that travelled to the medaka optic tectum despite the ectopic position of the retina (Figure 3B, C). The optic nerve from one ectopic retina usually innervated both *ipsi* and *contralateral* optic tecta (Figure 3C). The premature birth of zebrafish RGCs guarantees that their projections arrive to the target tissue earlier than the endogenous optic nerve, which results in a massive innervation of the medaka tecta by the zebrafish optic nerve (Figure 3B, C). Following the same rationale, the medaka optic nerves in medrakas arrive to the zebrafish tecta later that the endogenous nerve, and therefore its ramification is very limited (Supplementary Figure 4). These observations reveal an hetero-chronic formation of the same cell type within a chimeric embryo, following a genetic timing of differentiation despite sharing the physiological domain.

We observed examples in which the earlier formation of zebrafish RGCs resulted in anomalous optic nerve projections in zebrakas, evidencing that the optic nerve can indeed highjack on ectopic projection paths that are present at the time of navigation. That was the case for three zebrakas, in which the RGCs projected along the lateral line nerve (Figure 3D), which is present in the embryo before the pathfinding cues to reach the optic tecta. These miss-projections usually reached the caudal fin, evidencing a promiscuous behaviour of the zebrafish optic nerve in medaka hosts. Since these chimeras all projected to the optic tecta as well (Figure 3D, asterisk), our interpretation is that the earlier development of the lateral line nerve might have offered a permissive migratory route. Our results indicate that even when medaka and zebrafish blastocysts do not intermingle during epiboly and axis formation, differentiated cells can later on recognise cues present in the host, as those necessary for axonal pathfinding.

### Different sources for lens recruitment in zebraka and medrafish

The vertebrate eye is composed of the neuroepithelial derivatives, i.e. the neural retina (NR) and the retinal pigmented epithelium (RPE), which differentiate from a common progenitor pool [9,29] [30], and additional tissues in the anterior segment of the eye that derive from different germ layers [31]. The lens, a distinctive feature of the vertebrate eye, derives from the surface ectoderm and its formation is induced by retinal progenitors during early retinogenesis [31]. Lens induction therefore constituted yet another paradigm to assess inter-relations between donor and host tissues in chimeric embryos. When analysing the transcriptomes of zebrakas, we noticed that although retinal genes were represented (Figure 2E), lens transcripts from the donor were absent in the chimeras (Figure 4A). This indicate that the lens that is evident in zebraka either expresses a different set of transcripts or alternatively, it was formed using cells from the host.

**Figure 4.**
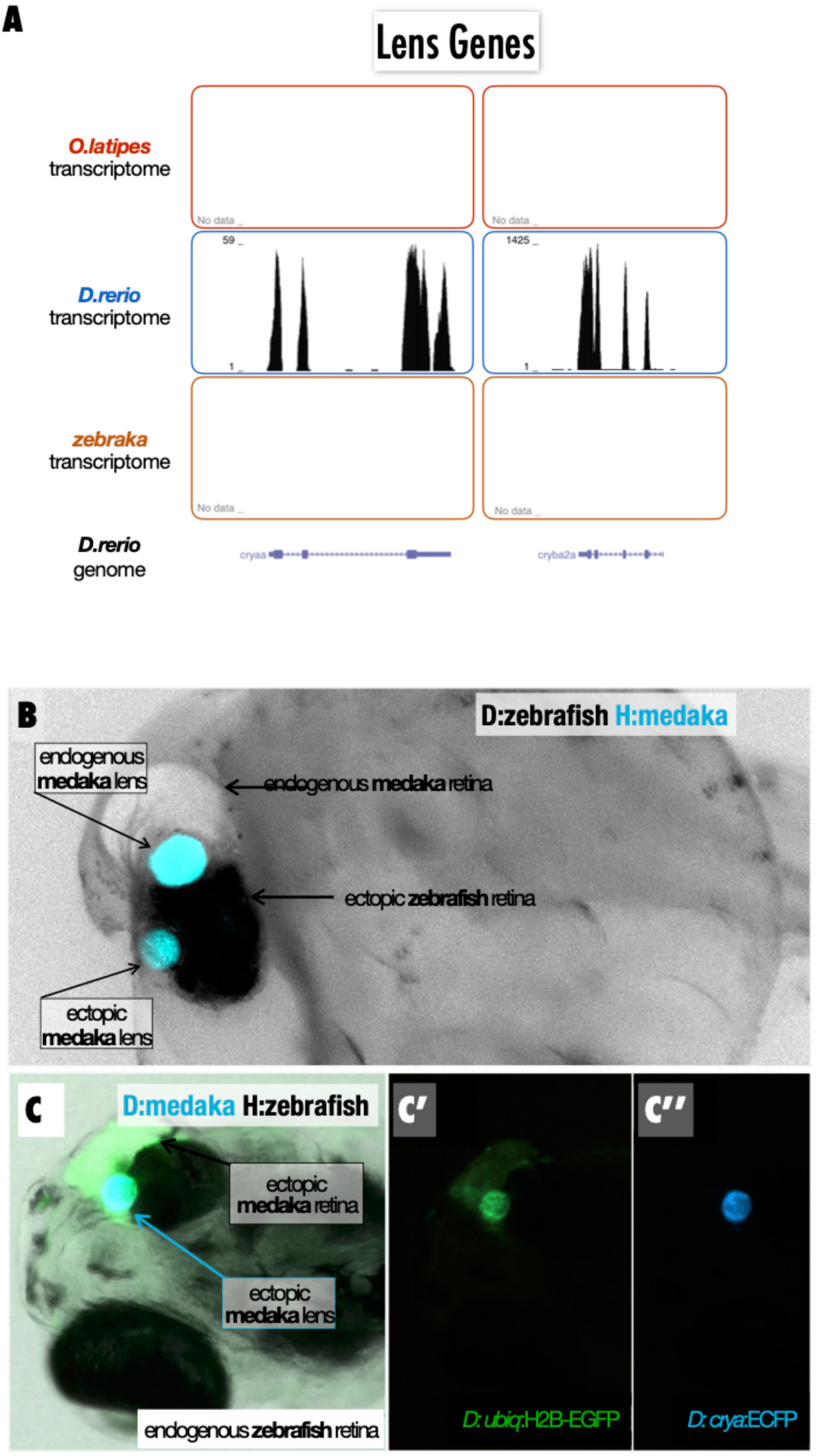
Ectopic zebrafish retina in zebrakas recruits the lens from the medaka host. **(A)** Transcriptomes of medaka (upper), zebrafish (middle) and zebraka (bottom) plotted along the zebrafish genome. Lens zebrafish genes (*crystalin a* and *crystalin b a2a, cryaa* and *cryba2a* respectively) are expressed in zebrafish but not in zebrakas, despite the fact that zebraka ectopic retina does contain a lens. **(B)** Fluorescent binocular picture (negative) of a zebraka showing the recruitment of an endogenous lens (host is medaka *crya*:ECFP) by an ectopic zebrafish retina, Tg (*OlBactin2*:EGFP). **(C)** Fluorescent binocular picture of a medrafish showing the ectopic medaka retina (EGFP+) contains also a medaka lens (donor is medaka *crya*:ECFP). Note that the endogenous zebrafish lenses do not express any fluorescent protein.

We performed blastula transplantations using EGFP+ blastomeres from zebrafish into the *cryA*:eCFP line from medaka, which labels the endogenous lenses with a cyan fluorescent protein [12]. When generated using this set up, zebrakas display an ectopic zebrafish retina which lens expresses the medaka *cry*:EGFP transgene (Figure 4B). This clearly indicates that zebrafish retinal cells recruited medaka host cells to form an ectopic lens. In the context of our previous observations showing that in zebrakas the ectopic retina develops earlier than the host retinae, our results evidence that the surface ectoderm has the potential to become a lens before the stage at which it is recruited endogenously in medaka. Surprisingly, this situation differs in the case of medrafish. When we transplant blastomeres from a medaka *cryA*:ECGF, *Gaudi*^LoxPOUT^ into unlabelled zebrafish host, we noticed that the ectopic retina displays a cyan lens (Figure 4C-C’’). These results reveal that, in contrast to the case in zebrakas, the ectopic medaka neural retina does not induce lens formation from zebrafish surface ectoderm. Therefore, the EGFP+ medaka cluster in medrafish generates both the retina and anterior structures of the eye like the lens. It is therefore valid to speculate that by the time the medaka retina starts the lens induction program, the surface ectoderm in zebrafish is no longer competent to acquire lens identity. Temporal windows for inductive process have been reported in other systems like the Hensen’s node in chicken, and the Spemann-Mangold organiser in frogs [32-39].

Trans-species transplantations have been long used in developmental biology to address the most diverse subjects. The broad range of examples includes, but is not limited to, the pioneering quail-to-chicken experiments to follow lineages [40,41], graft transplantations from *Planaria dorotocephala* to *Planaria maculata* to demonstrate the potency of regenerating cells [42,43], grafting different plant species to address horizontal genome transfer and speciation [44], transplantation of blastomeres between related teleost to address primordial germ cell migration [45], rat to mice blastomere transplantation [46] to address the ‘empty niche’ hypothesis [47], human to mice cells for teratoma assays, and the recent human-to-pig chimeras to explore the boundaries of pluripotent stem cell transplantations in mammals [48]. Here, we developed a new experimental set-up for isochronic trans-species transplantations and used it to explore intrinsic and extrinsic temporal properties of retinogenesis.

We combined blastomeres of species that diverged 250 My ago and observed that although initially, cells grouped — or stayed grouped — in a species-specific fashion, later on the exogenous cluster interacts with the biochemical and/or physical environment of the host. Albeit not participating into the host morphogenesis, as previously reported for other inter-species transplantation in fish [45], the cluster expresses different retinal maker genes and generate retinal cell types. Moreover, it reads cues from the host to navigate axonal outgrowth, and displays inductive properties onto host cells as demonstrated by zebrafish cells inducing lens formation on the medaka host. Yet despite these interactions with the host, the ectopic retina followed a species-specific developmental time. Our experiments add to the growing evidence on autonomous organisation of biological systems [2,49,50] with a focus on developmental dynamics. It has recently been suggested that in mammals, developmental timing depends on protein stability, a kinetic parameter that has species-specific features [51]. Whether the same holds true for an entire extrinsic organ developing in a different host species is an attractive hypothesis that needs to be formally tested. Regardless of the molecular nature of the intrinsic timing, our results illustrate the temporal window in which a tissue can be induced into another (i.e., surface ectoderm into a lens), which proved to be way broader than the temporal requirements during embryonic development. Overall, our experiments report a successful chimerism between vertebrate species that differ more than 250 My ago, the highest distance for an isochronic transplantation in such an early embryonic stage. Using this novel inter-species paradigm to tackle temporal aspects of embryonic development, we could reveal some basic principles predicted by the modularity hypothesis [52]. Thus, here we show that the evolutionary conserved “retinal module” can be defined both by its intrinsic genetic identity and its external connectivity to neighboring modules.

## Acknowledgments

We are tankful to the Centanin and S. Lemke groups, J. Wittbrodt and A. Martinez Arias for scientific input on earlier version of the manuscript, J. Lohmann and T. Greb for feedback on plant inter-species transplantation experiments, N.Foulkes for constructive input regarding developmental dynamics, E. Leist, M. Majewski and A. Saraceno for fish husbandry. This work is supported by grants from the Deutsches Forschungsgemeinshaft to L.C. (SFB873 – A11) and the Spanish Ministry of Science, Innovation and Universities (MICINN) to J-R.M.M.: BFU2017-86339P and MDM-2016-0687. LB contract is supported by Fundación Ramón Areces-2016.

## Author Contributions

Conceptualization, LC; Methodology, JFF, LC; Software, LB, JRMM; Validation, JF; Formal Analysis, JFF, LB, JRMM, LC; Resources, JRMM, LC; Writing – Original Draft Preparation, LC; Writing – Review & Editing Preparation, JFF, JRMM, LC; Visualization, JFF, LB; Supervision, LC; Project Administration, LC; Funding Acquisition, JRMM, LC.

## Declaration of Interests

The authors declare no competing interests.

## Materials and Methods

### Fish stocks and transgenic fish lines

Medaka (*Oryzias latipes*) and Zebrafish (*Danio rerio*) stocks were maintained according to the local animal welfare standards (Tierschutzgesetz §11, Abs. 1, Nr. 1). Animal experiments were performed in accordance with European Union animal welfare guidelines (Tierschutzgesetz 111, Abs. 1, Nr. 1, Haltungserlaubnis AZ35–9185.64 and AZ35–9185.64/BH KIT).

The following zebrafish lines were used in this study: AB zebrafish strain as a wildtype, *OlBactin2:*EGFP-CAAX [7], *Atoh7*:GFP [15], *ef1a*:Lyn-Cherry

The following medaka lines were used in this study: Cab strain as a wild type, Heino (Albino medaka) (reference), Wimbledon DsTrap#6 [7], *Rx2*::H2B-mRFP [13], *zFli1*::EGFP [53], Gaudi^RSG^ (contains the integration reporter *crya*:ECFP that drives ECFP expression in the lens), Gaudi^LoxPOUT^ [12], *Atoh7*::EGFP [15], *Atoh7*::lynTdTomato [54].

### Intra- and Inter-species transplantation

Zebrafish crosses were set up at 10 a.m, collected after 20 min and kept at room temperature. Medaka couples were maintained together and produced from 8:00 am on. Medaka eggs were collected between 9.00 and 10.30 a.m. and grown at 32°C to synchronize embryos at blastula stage. Blastula stage embryos were dechorionated as previously described using hatching enzyme for medaka and pronase (30mg/ml) for zebrafish, and placed for transplantation in agarose wells with the proper medium for the host species (E3 medium for zebrafish or ERM for medaka).

One blastula could be used as a donor for 3-5 hosts, 20 to 50 cells were transplanted from the animal pole region of the donor to the hosts animal pole. Transplantations were carried out as previously described [55]. Transplanted embryos were kept in growth medium of the host species. In accordance with animal welfare standards transplantations with Zebrafish hosts were maintained up to day 5 of embryonic development and Medaka hosts were grown to day 9 of embryonic development.

### Antibodies and staining

Primary antibodies used in this study were rabbit anti-GFP (Live technologies, 1/750), chicken anti-GFP (Invitrogen, 1/750), rabbit α-DsRed (1/500). Secondary antibodies were Alexa488 anti-rabbit (Live Technologies 1/500) and Alexa488 anti-chicken and Dylight 548 anti-rabbit (Jackson 1/500). DAPI was used in a final concentration of 5 ug/l.

### Imaging

Stained embryos were imaged with a laser-scanning confocal microscope Leica TCS SP8 (20x immersion objective) or a Leica TCS SPE. Imaging was done on glass-bottomed dishes (MatTek Corporation, Ashland, MA 01721, USA).

Live embryos were anaesthetized in 1mg/ml Tricaine in the respective fish medium as previously described [56] and imaged in 3% Methylcellulose in ERM or in 0.6% low melting agarose in ERM. Embryos were screened and imaged using a Stereomicroscope (Olympus MVX10 MacroView) coupled to a Leica DFC500 camera or at a laser-scanning confocal microscope Leica TCS SP8 (20x immersion objective) or Leica TCS SPE. All subsequent image analysis was performed using Fiji software [57].

### RNA seq on Zebrakas

Zebrakas with a EGFP+ cluster in the head at 50hfp were used to extract total RNA (Trizol), together with zebrafish donor and medaka hosts that were grown at the same temperature. Libraries were prepared from total RNA followed by a polyA selection (NEBnext PolyA) and sequenced in a NextSeq 500 platform. They produced an average of 69M 85-nt single end reads for each sample. RNA-seq samples were mapped against both oryLat2 and danRer10 assemblies using Hisat2 [58]. Datasets can be accessed at GEO: *waiting for confirmation from the repository*. The aligned read SAM files were assembled into transcripts and their abundance was estimated using Cufflinks v2.2.1 [59]. Zebraka FPKM ratios between retinal developmental progression genes (*atoh7, ptf1a* and *pou4f2*) and retinal progenitor markers (*rx3* and *vsx2*) were compared with the same ratios from wt medaka and zebrafish transcriptomes at resembling developmental stages [26]. In the case where both the numerator and the denominator of the ratio were equal to zero, the resulting value was also reported as zero.

## Supplementary Figure Legends

**Supplementary Figure 1.**
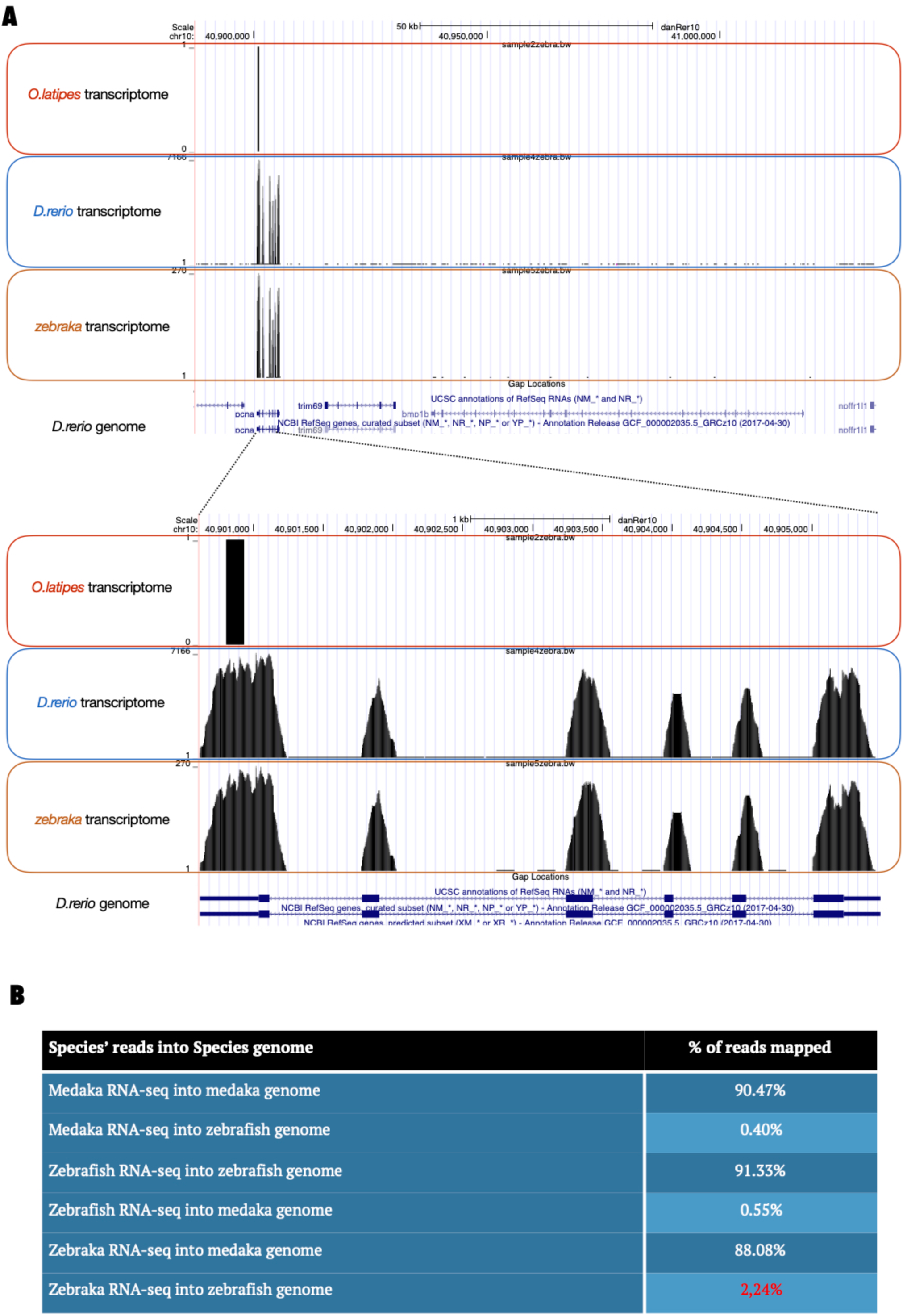
Transcriptomes of medaka, zebrafish and zebraka plotted on the zebrafish genome. **(A, Top)** Transcriptomes from medaka are in the upper compartment, from zebrafish in the middle and from zebraka at the bottom. The image shows an example of a peak obtained using the medaka transcriptome. **(A, Bottom)** A detail of the region shows a different morphology for the medaka peak, different from the zebrafish or zebraka peaks that span along the entire set of exons of PCNA. Also note that the medaka peak consist of just one read, different from the >7K in zebrafish or >250 in zebraka. **(B)** Table displaying the values of reads mapped using transcriptomes and genomes from both species.

**Supplementary Figure 2.**
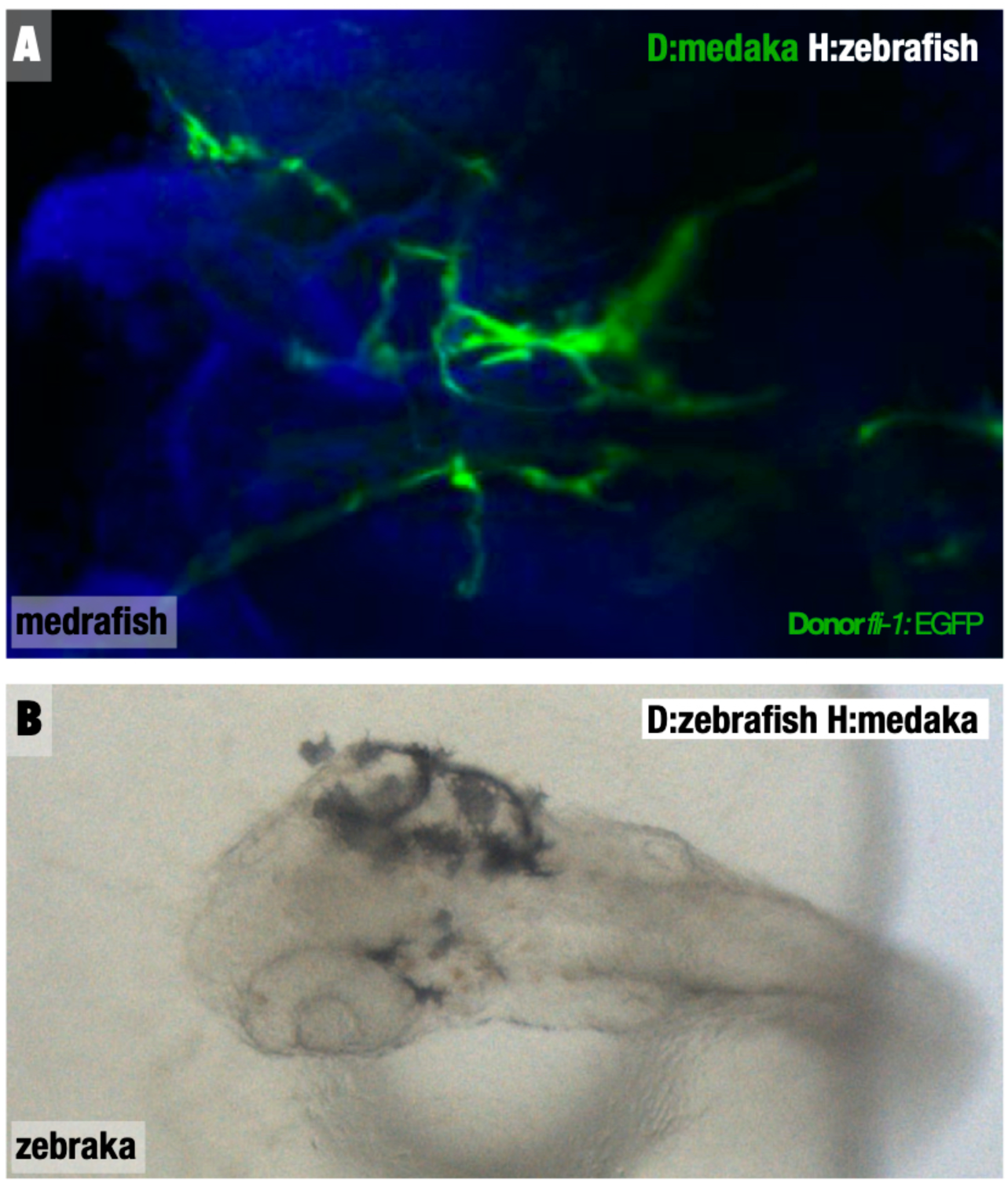
Differentiation of donor cells in inter-species transplantations. **(A)** Confocal image of a 3 dpf medrafish using Tg(*fli1*:EGFP) medaka as a donor. **(B)** Binocular picture of a 2.5 dpf zebraka using a pigmented zebrafish as a donor and an albino medaka as a host.

**Supplementary Figure 3.**
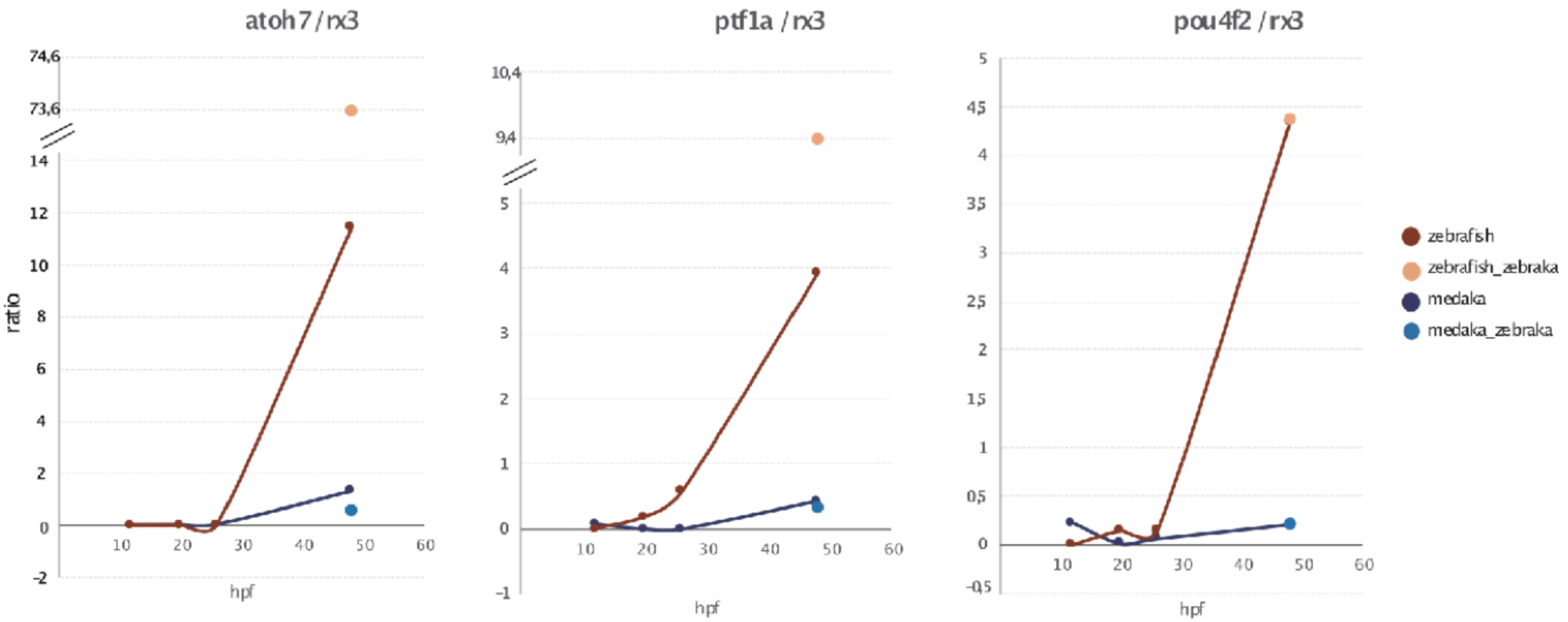
Ratios of progenitor / differentiation retinal genes in zebrakas. Plots show lines for the specified ratio during early embryonic development in zebrafish and medaka. The dots at 50hpf correspond to samples used in this study.

**Supplementary Figure 4.**
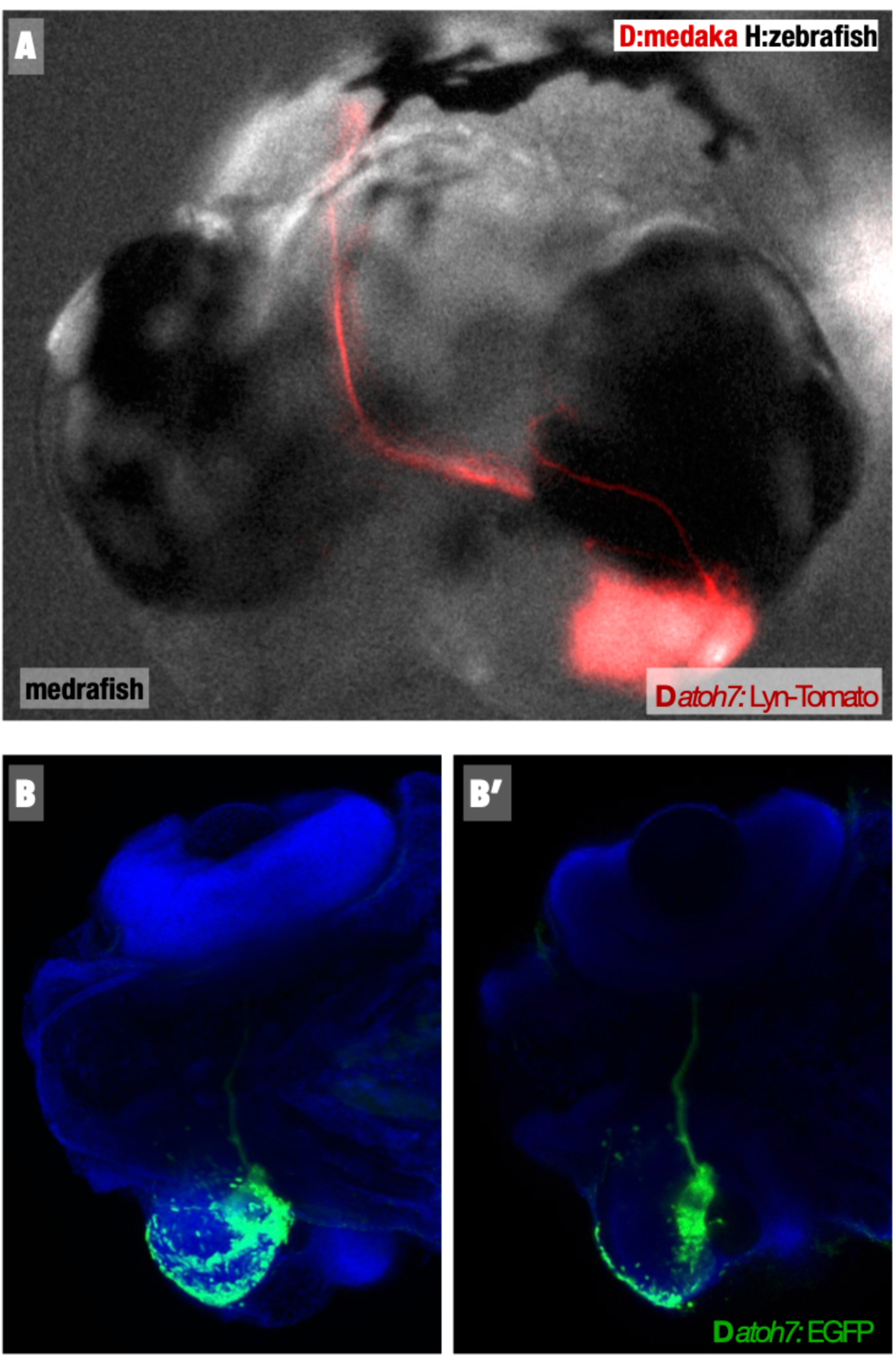
Differentiation of RGCs is an ectopic medrafish retina. Confocal image of a 4 dpf medrafish using Tg(*atoh7*:LynTomato) medaka as a donor and a non-transgenic zebrafish host. RGCs differentiate in the ectopic retina and form an optic nerve that projects to the host tectum.

